# Cell-type specific allelic dampening of sex-linked genes in sex chromosome aneuploidy

**DOI:** 10.64898/2026.04.16.719032

**Authors:** Gala N. Filippova, Elliott Sanger, James MacDonald, He Fang, Camille Groneck, Masaya Takasaki, Anastasia Meleshko, Wenxiu Ma, Yajuan Liu, Gang Li, Ran Zhang, Charles E. Murry, Daniel Van Dyke, Anne Skakkebæk, Claus H. Gravholt, William S. Noble, Theo Bammler, Jessica E. Young, Xinxian Deng, Joel B. Berletch, Christine M. Disteche

## Abstract

Common sex chromosome aneuploidies (SCAs) often present with cognitive and cardiovascular dysfunction in humans. To address SCA effects on gene expression and DNA methylation in relevant cell types, we differentiated neural precursor cells (NPCs) and cardiomyocytes (CMs) from human induced pluripotent stem cells (hiPSCs) with different numbers of sex chromosomes, including isogenic and independent lines. As expected, the expression of genes that escape X inactivation (escapees) mostly increases with the number of inactive X chromosomes (Xi). However, allelic analysis shows dampening of escapees specifically on the Xi in XXY compared to XX in both NPCs and CMs, revealing a novel type of dosage compensation in SCA. In contrast, Y-linked gene expression is higher in XXY versus XY NPCs, but the opposite is observed in CMs. This may explain the greater number of differentially expressed autosomal genes in NPCs versus CMs with an added Y chromosome, while effects of added X chromosomes are similar between cell types. Concordantly, changes in autosomal DNA methylation are mainly driven by the presence of a Y chromosome. These findings highlight the cell-type specificity of sex-linked and autosomal gene regulation in SCA conditions.

**Highlights:**

- Sex chromosome aneuploidy induces cell-type specific changes in gene expression
- Dampening of the inactive X chromosome in XXY alleviate X overexpression
- High Y-linked gene expression in XXY neuronal precursor cells but not cardiomyocytes
- Sex chromosome aneuploidy disrupts sex biases in autosomal gene expression

## Introduction

Sex chromosome aneuploidies (SCAs), such as Klinefelter syndrome (XXY), Triple X syndrome (XXX) and Turner syndrome (X), often cause behavioral and cardiac disorders in humans^[1–3]^. Although SCAs with two or more X chromosomes result in inactivation of all X copies (Xi’s) but one (Xa), a number of genes called “escapees” remain expressed from the Xi. These genes are under-expressed in Turner syndrome and over-expressed in the presence of supernumerary X chromosomes. Cumulative effects of abnormal gene dosage in SCAs contribute to phenotypes such as behavioral abnormalities, a lower IQ, and cardiac disorders. Anatomical abnormalities including abnormal size of the brain and that of specific brain regions track with the number of sex chromosomes^[4]^. Similarly, cardiac abnormalities can have opposite features dependent on the type of SCA, for example, a prolonged heart-rate corrected QT (QTc) interval in Turner syndrome and a shortened one in Klinefelter syndrome^[5]^.

SCA anomalies are due to a combination of hormonal effects and direct and indirect effects of abnormal dosage of sex-linked genes^[6–9]^. These genes belong to one of the following categories: (1) pseudoautosomal (PAR) genes expressed from both X and Y chromosomes, albeit at a slightly reduced level from the Xi compared to that from the Xa or the Y; (2) non-PAR escapees, some retaining paralogs on the Y chromosome (X/Y genes); (3) X-linked genes transcriptionally silenced by X chromosome inactivation (XCI); and (4) Y-linked genes unique to individuals with at least one Y chromosome. Importantly, the number of escapees and their expression levels from the Xi versus the Xa can vary among cell types, stressing the need for allele-specific and cell-type-specific analyses of SCAs^[10–12]^. Few specific sex-linked genes have been implicated in SCA phenotypes in part due to the lack of accessible relevant tissues and cell types. Few specific sex-linked genes have been implicated in SCA phenotypes in part due to the lack of accessible relevant tissues and cell types. Two examples are *ZFX* implicated in abnormal brain anatomy^[9]^ and *SLC25A6* implicated in cardiac anomalies^[5, 13, 14]^. SCAs are associated not only with abnormal expression of sex-linked genes but also of autosomal genes, and affected genes include protein coding genes, as well as genes encoding microRNAs, lncRNAs, or circular RNAs^[15, 16]^. DNA methylation changes are also associated with specific SCAs, for example, the genome is hypomethylated in Turner syndrome and hypermethylated in Klinefelter syndrome^[7, 15, 16]^. However, specific correlations between gene dysregulation and DNA methylation are limited.

Most analyses of SCA effects on gene expression and DNA methylation have been carried out in cell lines such as lymphoblastoid and fibroblast lines, and in easily obtained tissues such as blood, with few studies focusing on other tissues such as fat and muscle^[7, 17, 18]^. Mouse models of SCAs in which *in vivo* tissue and cell type-specific analyses can be performed have the advantage that littermates provide a controlled isogenic system^[19–22]^. However, SCA phenotypes are much milder in mouse than human, owing to the presence of fewer escapees in mouse and to functional differences of sex-linked genes^[23]^. Thus, approaches to examine cell types relevant to SCA phenotypes, such as neural cells and cardiomyocytes, in a controlled genetic background are essential to better understand SCA effects on human phenotypes. Here, we derived pairs of isogenic human induced pluripotent cell (hiPSC) lines with different numbers of sex chromosomes. Following differentiation into neural precursor cells (NPCs) and cardiomyocytes (CMs), paired isogenic cell lines, together with independent lines, were analyzed by RNA-seq, including allele-specific analysis, and by DNA methylation analysis.

## Methods

### hiPSC reprogramming and generation of isogenic pairs

hiPSC lines X4, X5, and X6 were reprogrammed from primary amniotic fluid-derived XXY cell lines GM02269, GM03091, and GM03535, respectively (from the Coriell Institute). hiPSC line X3 was derived from a mosaic X/XXX cell culture grown from products of conception (X3: mos X/XXX). Normal controls hiPSC lines (male lines M7, M8, M9 and female lines F9, F11, F12) were reprogrammed from independent amniotic fluid-derived cell lines with normal XY or XX karyotypes, respectively. Reprogramming of amniotic fluid-derived and products of conception-derived cell lines was done using non-integrating episomal factors (Addgene: 27077, 27078, and 27080) and standard protocols^[24]^. hiPSC lines X175 and X183 were reprogrammed from mosaic XY/XXY peripheral blood mononuclear cells (PBMCs) obtained from adults with Klinefelter syndrome, using a non-integrating Sendai viral kit (Cytotune 2.0 kit from Thermofisher). hiPSCs were reprogrammed and initially cultured on inactivated mouse embryonic fibroblasts (MEFs) and pluripotency was confirmed by morphology and expression of known pluripotency markers (Figure S1A, B). The hiPSC lines were further differentiated to embryoid bodies (EBs) that were positive for expression of all three germ layer markers as previously described^[25]^. All original material was de-identified so that this study was considered non-human subjects research by the University of Washington Institutional Review Board.

Generation of isogenic pairs of hiPSC lines with a different number of sex chromosomes was achieved using two methods (Figure S1C). First, we derived an isogenic XY line from the XXY hiPSC line X5. Using AAV-mediated homology-based recombination, a 2.3 kb TK (thymidine kinase)-neomycin (neo) dual-selection cassette (driven by the EF1a promoter and flanked by *XIST* homology arms) was inserted into the *XIST* intron 1 region (insertion location: chrX:73839709, hg38). Following selection using G418 as described^[26]^, integration of the TK-neo cassette in *XIST* was confirmed by PCR and sequencing using primers flanking the conjunction sites. ∼30% of G418-resistant clones had the TK-neo cassette integrated into the *XIST* locus. Note that this integration could occur on the Xi, the Xa, or both in any given cell. G418-resistant clones with the correct integration were then subject to negative selection using ganciclovir against cells expressing TK, resulting in viable cells with spontaneous loss of the Xi or with TK null mutations or epigenetic silencing of the cassette. To identify isogenic clones with loss of the Xi, ganciclovir-resistant single-cell colonies were expanded and screened by RT-PCR for loss of *XIST* expression (Figure S1D). The genotypes of the isogenic clones were further verified by *XIST* copy number qPCR (Figure S1E) and DNA-FISH with an X-specific probe (Figure S1F). An alternative approach to obtain isogenic pairs was to isolate single-cell clones of hiPSCs derived from XY/XXY mosaic cultures X175 and X183 and from X/XXX mosaic culture X3 (Figure S1C). As described above, to identify isogenic clones of each genotype, all single-cell derived hiPSC clones were screened for *XIST* expression, followed by *XIST* copy number qPCR and karyotyping or DNA-FISH to confirm the genotypes, which were further verified based on DNA methylation patterns determined by array analysis (see below and Figure S2C, D). Prior to differentiation, all hiPSC clones were adapted to feeder-free conditions, and lack of episomal vector integration as well as clearance of Sendai virus were confirmed by RT-PCR according to the manufacturer’s protocols. Most of the iPSCs were clear from Sendai expression by passage 5-8. All hiPSC clones with more than one X chromosome (XX, XXY, and XXX) were also routinely frozen at early passages and tested for robust *XIST* expression by qRT-PCR to exclude clones with the erosion of XCI. Only clones with stable XCI (the highest *XIST* expression) and at the earliest passages were used for differentiation. See Table S1 for additional details on the cell lines.

### NPC differentiation

hiPSCs were differentiated into NPCs using a protocol adapted from previous studies (Figure S1G)^[27–31]^. Briefly, cells were densely plated (1-1.5 x 10^6^ per well) into 12-well plates pre-coated with laminin-521 (BioLamina). NPC differentiation was induced in neural Induction Media (NIM) consisting of BNMM media [(1:1 mix of DMEM/F12 (Gibco) and neural basal media (Gibco) supplemented with N2 (Gibco), B27 (Gibco), non-essential amino acids (Gibco), Glutamax-100 (Gibco), insulin-transferrin-selenium-sodium pyruvate (ITSA) (Gibco), beta-mercaptoethanol (Gibco), 1% penicillin-streptomycin (Gibco)] supplemented with dual SMAD inhibitors SB431542 (10μM) and LDN193189 (0.5μM) (Biogems). Following 9 days of neural induction, cells were treated with Versene (Gibco) and collected using a cell scraper. After gentle trituration, cells were transferred to 12-well plates pre-coated with laminin-521 and fed with BNMM (without SMAD inhibitors) for 3 days. After emergence of neural rosettes, cells were fed with BNMM media supplemented with 20 ng/ml FGF (Fisher Scientific) (Figure S1G, top). Cells were collected for molecular analyses at day 19 of differentiation. Differentiation to NPCs was confirmed by morphology and immunostaining for nestin (Santa Cruz #sc-23927). Briefly, cells fixed for 30 min in 4% paraformaldehyde were washed in PBS and blocked in PBS containing 2.5% BSA and 0.1% Triton X 100 for 30 min at room temperature (RT) prior to applying a primary nestin antibody at a 1:300 dilution and incubation at 4°C overnight. A secondary antibody (Invitrogen #A11032), was then applied at a 1:500 dilution for 1 h at RT. Cells were counterstained using DAPI and stored in Vectashield anti-fade before imaging on a high-resolution wide-field fluorescent microscope (Figure S1H).

### CM differentiation

hiPSCs were differentiated into CMs using published protocols with minor modifications (Figure S1G)^[32, 33]^. Briefly, cells were seeded at density 2-3 x 10^5^ per well into Matrigel (Corning) coated 12-well plates in mTeSR+ media (Stem Cell Technologies) supplemented with ROCK inhibitor 10μM Y-27632 (Selleckchem). After 24 h, the media was changed to mTeSR+ with 1μM CHIR-99021 (Stemgent). On the following day (day 0 of differentiation), the cells were switched to RBA media [RPMI with 500 μg/ml BSA (Gibco) and 213 μg/ml ascorbic acid (Sigma)] supplemented with 4μM CHIR-99021. On day 2 (48 h later), the media was replaced with RBA supplemented with 2μM of the WNT inhibitor Wnt-C59 (Selleckchem). On day 4, the cells were fed with RBA (with no small molecules added) and incubated for 2 more days. Finally, on day 6 the cells were switched to RPMI media supplemented with B27 (Gibco), with feedings occurring every other day until collection on day 21. Initial beating was usually observed between days 7-11. Differentiation of CMs was confirmed by morphology and flow cytometry using a cardiac troponin T (cTnT) antibody (Invitrogen) (Figure S1G, bottom and S1H). Cardiomyocytes were dissociated using warm Versene (Gibco) supplemented with 0.5% trypsin for 5 min at 37° and collected in Stop solution [RPMI supplemented with 10% fetal bovine serum (FBS)] to neutralize the trypsin. Cells were spun down and fixed in 300 μl of cold 4% paraformadehyde for 10 min at RT. After fixation, the cells were resuspended in 300 μl PBS + 5% FBS and split into two wells of a 96 well plate for staining with either a mouse cTnT antibody (1:100) (Thermofisher MS-295) or a mouse control isotype IgG1 (1:100) (eBioscience 14-4714) for 30 min at RT. After washing, both sets were incubated with a goat anti-mouse phycoerythrin (PE) secondary antibody (1:200) (Jackson 115-116-072) for 30 min at RT in the dark. All stainings and washes were performed in the presence of 0.75% saponin (Sigma) in PBS + 5% FBS. For flow cytometry analysis, cells were washed and resuspended in PBS + 5%FBS and run on FACSCanto II (BD Biosciences). FloJo was used for the analysis. Only cells > 80% cTnT positive were used for downstream analyses (Figure S1I).

### RNA-seq analysis

NPCs and CMs were collected and stored in Qiazol (Qiagen) prior to RNA isolation. For all cell lines, bulk RNA-seq indexed libraries were prepared using Illumina TruSeq RNA kit V2 from 1µg of total RNA. Sequencing was done on a NextSeq sequencer to yield 75bp single-end reads. FASTQ files were aligned to the GRCh38 genome using the STAR aligner^[34]^, using a conventional genome index for male samples, and a genome index generated using a masked Y chromosome (i.e. all bases converted to N) for female samples, to eliminate cross-alignments to the PAR regions on the Y chromosome. Read counts per gene were generated from BAM files using the Bioconductor Rsubread package^[35]^. For hiPSCs, gene expression was estimated using Tophat/v2.0.14^[36]^ with default parameters and gene-level expression was normalized using FPKM (fragments per kilobase of exon per million mapped reads). Differential expression was determined using cuffdiff^[37]^. For analysis in differentiated cells, surrogate variable analysis (SVA) was used to remove batch effects prior to downstream analyses^[38]^. To identify differentially expressed genes (DEGs) in NPCs and CMs pairwise comparisons of cell lines were set up as detailed in Table S2. The Bioconductor edgeR package was used to analyze the data, using the limma-voom pipeline that converts counts to log counts/million and then fits a weighted linear model to account for heteroskedasticity^[35]^. Models were fit with blocking by subject when possible. Otherwise, within-subject correlations were estimated using the duplicateCorrelation function, and a generalized least squares (GLS) model was used to account for the correlation structure. Genes with an unadjusted p<0.001 were considered significantly differentially expressed.

### DNA methylation analysis

DNA isolated from NPCs and CMs was subjected to bisulfite conversion according to the manufacturer’s protocol (Zymo). DNA methylation data was then generated using Infinium MethylationEPIC BeadChip (850k) (Illumina). Prior to data analysis, cross-reactive probes, probes with SNPs proximal to the CpG sites, and probes that were indistinguishable from background binding were excluded. The Bioconductor minfi package was used for preprocessing and normalization by a functional procedure that uses “out of bounds” probes to estimate background and control probes to remove technical variation^[39–41]^. In addition, an X/Y interpolation from the Bioconductor wateRmelon package^[42]^ was used to normalize CpGs found on the allosomes. The beta estimates (proportion of methylation at each CpG site) were converted to M-values using a logit transform for all further analyses. The Bioconductor limma package was used to make comparisons between groups at the CpG level^[43]^, blocking on subject when possible, and using duplicateCorrelation to estimate within-subject correlation, which was then used to fit a GLS (Generalized least squares) model that accounts for within-subject correlations when blocking on subject is not possible. Individual CpG sites were considered differentially methylated positions (DMPs) at a false discovery rate (FDR<0.05)^[44]^. Differentially methylated regions (DMRs) were identified using the Bioconductor DMRcate package^[45]^, which uses limma to estimate the CpG-level differences between groups and then identifies DMRs using a Gaussian kernel function, based on the defaults of a 1000 bp kernel support and ≥2 CpGs in a DMR. Note that FDR<0.1 was used to identify CpGs within DMRs in DMRcate.

### X-linked allelic expression analysis

Allelic analysis of X-linked gene expression was done on each clone of XX and XXY NPCs and CMs, in which single-cell cloning resulted in completely skewed XCI. Only reads uniquely mapped to the X chromosome were retained for compiling SNP read counts using cellsnp-lite with a minimum read count (minCOUNT) set to 5, minimum minor allele frequency (minMAF) set to 0.05, and minimum mapping quality score (minMAPQ) set to 20^[45]^. Allele frequency information was sourced from the cellsnp-lite documentation [https://cellsnp-lite.readthedocs.io/en/latest/main/data.html#data-list-of-common-snps], which contains 36.6 million SNPs called from the 1000 Genomes Project^[46]^ with a minor allele frequency (MAF) > 0.0005 in hg38. To derive allele-specific counts at each SNP, we performed the following steps using scripts sourced from the scLinaX preprocessing vignette ^[47]^ (https://ytomofuji.github.io/scLinaX/articles/scLinaX_preprocessing_example.html): Raw SNP read count matrices generated by cellsnp-lite were processed to extract allele-specific counts for each clone. Alternative allele read counts (ALTcount), total read depth (DP), and other allele read counts (OTHcount) were loaded, and reference allele counts (REFcount) were derived as DP – ALTcount, consistent with the direct read pileup against the reference base. SNPs were annotated with their genomic coordinates and reference (REF)/alternative (ALT) alleles from the associated VCF file. The processed data was reformatted into a long-format table where each row represents a unique SNP-clone pair, containing columns for the gene name, SNP position, XCI status, SNP mean proportions, individual donor SNP proportions, and statistics (Additional dataset 1). The SNP read count data was further filtered based on functional annotations obtained from ANNOVAR^[47]^, and SNPs were matched to their corresponding genomic annotations, retaining only those located in exonic, intronic, UTR, ncRNA, or splicing regions of a gene. Variants mapping to multiple genes or lacking gene annotations were removed, and the final QC-passed SNP dataset was saved for downstream analysis. SNP read counts were summed across technical replicates (denoted by rep1, rep2). In all further analyses, technical replicates were treated as one clone. Finally, annotated SNP dataframes from all clones and cell types were concatenated into a single, comprehensive dataframe for downstream analyses, and each gene was labeled with their XCI status sourced from^[47]^. In total, there were 14,039 SNPs from 30 clones across eight individuals and two cell types.

To estimate the proportion of escapees and compare escape patterns across genotypes, an analytical pipeline was implemented as follows. First, to reduce the effects of sequencing errors, an error proportion for each SNP was calculated by dividing the lowest read count among REF, ALT, and OTH allele read counts by the total read count across these alleles. SNPs with an error proportion greater than 0.01 were excluded from further analysis. Next, the SNP-level XCI proportion was determined by assuming that for each SNP, the allele with the highest read count corresponds to the Xa, while the allele with the second highest read count corresponds to the Xi. The proportion of Xi reads was calculated as Xi / (Xa + Xi) for each SNP. An exception was made for *XIST*, where the allele with the highest read count corresponds to the Xi. Additionally, SNPs with a total read count (Xa + Xi) below the minimum read count threshold of 20 were excluded. To ensure a robust approximation of the Xi proportion, each gene’s Xi proportion was calculated as the Xi proportion of the SNP with the highest total read count (Xa + Xi). Subsequently, Xi proportions were averaged across clones from the same individual donor to derive an individual-specific Xi proportion. Considering genes with data from at least three individual donors per genotype, differences in Xi proportions between genotypes were tested using a two-sample t-test using the ttest_ind function from the scipy^[48]^ package with default parameters. The resulting p-values were then adjusted for multiple comparisons using the Benjamini-Hochberg correction, implemented via the multipletests function from the statsmodels package^[49]^. All results were compiled into a dataframe (Additional data set 1). Finally, gene locations were annotated based on Release 47 (GRCh38.p14) from GENCODE^[50]^. Since the genomic sequences of individual donors or their parents are not available, our allelic analysis of X-linked genes in cloned XX and XXY lines with skewed XCI is limited to X-linked genes that have some level of biallelic expression of SNPs in a given cell line.

Clones derived from each XX or XXY individual donor may differ in terms of which X chromosome (maternal or paternal) is inactivated due to random XCI. To identify the Xi in each clone, we used allele-specific expression profiles of SNPs, excluding those within the PAR genes and filtering SNPs to retain only those with a total read count above 20. To remove sequencing and mapping noise, an error proportion threshold >0.01 was used as described above. For each SNP that passed filtering, the allele with the lowest count was zeroed out, under the assumption that it likely reflects noise or sequencing error. The remaining two allele counts (the major and minor) were normalized so their sum equals 1, producing per-allele proportions that represent relative Xa or Xi expression. Next, pairwise comparisons were done using only SNPs shared between samples. For each shared SNP, the normalized REF, ALT, and OTH proportions from one sample were paired with those from the other sample. Pearson correlation coefficients were computed separately for the REF, ALT, and OTH allele proportions across the shared SNPs. An “Overall correlation” was computed by concatenating all REF, ALT, and OTH proportions into a single vector per sample and calculating the Pearson correlation between the two vectors. SNPs where both samples had zero expression for a given allele were excluded from the correlation calculation for that allele (Table S5).

## Results

### Generation of NPCs and CMs with differential sex chromosome content

To evaluate the consequences of SCA on gene expression and epigenetic changes genome-wide in relevant cell types and to reduce the effects of individual variation, we derived isogenic hiPSC lines with different numbers of sex chromosomes in a controlled genetic background for subsequent differentiation in NPCs and CMs (Table S1). Two pairs of XY/XXY and one pair of X/XXX isogenic hiPSC lines were derived from mosaic individuals, with one additional XY/XXY pair derived from an XXY hiPSC line in which loss of the Xi was selected by insertion of a selectable marker at *XIST* (see Methods) (Figure S1C)^[26]^. Independent hiPSC lines were derived from three XXY individuals as well as three control XX female and three control XY male individuals. Once verified by karyotyping, single-cell derived hiPSC clones were differentiated to NPCs or CMs (See Methods, Figure S1G and Table S1). Note that the X line from the X/XXX pair was not differentiated into CMs. NPC and CM differentiation of each hiPSC clone was confirmed using cell morphology and immunostaining of cell type-specific markers (nestin for NPCs and cardiac troponin T for CMs) (Figure S1H, I). hiPSCs with more than one X chromosome and their derivatives are prone to XCI erosion via loss of *XIST* expression and DNA methylation, which can lead to aberrant reactivation of the Xi^[51, 52]^. Thus, prior to downstream analyses, robust *XIST* expression was confirmed in each XX, XXY, and XXX hiPSC clone prior to- and post-differentiation to NPCs or CMs (See Methods and Figure S2A, B). We observed that stable XCI was maintained during differentiation of these clones to NPCs and CMs, which was further verified based on DNA hypermethylation at CpG islands of X-linked genes (Figure S2C, D). A summary of lines/clones can be found in Tables S1 and S2.

### Sex-linked gene expression in SCA

The effects of SCA on expression of X- and Y-linked genes, which could be either direct or indirect, were determined in NPCs and CMs by systematic identification of differentially expressed genes (DEGs) between XXY and XX or XY lines, and between XX and XY lines, using both isogenic pairs of XXY/XY lines and independent XXY, XY, and XX lines (Figure 1A, B; Table S2). Note that a single XXX/X isogenic pair was included in overall comparisons of gene expression but excluded in comparisons of individual genes due to sparsity of data. Overall, hierarchical clustering shows a clear separation of NPC and CM clones based on the number of X’s and the presence of a Y (Figure S2E, F). X- and Y-linked gene expression was found to be less variable between partners of isogenic pairs than among independent lines, reflecting mitigated genetic variability in isogenic pairs (Figures 1A, B and S3A, B). As expected, expression of PAR1 genes consistently increases with the total number of sex chromosomes in both NPCs and CMs, with the highest expression observed in cells with a Y, reflecting the known partial repression of PAR1 genes on the Xi (Figures 1A, B and S3C, D; Table S3)^[12]^. In contrast, the majority of non-PAR escapees^[11, 53]^ have higher expression in lines with more than one X chromosome in both NPCs and CMs, reflecting cumulative expression from the Xa and Xi (Figures 1A, B and S3A, B; Table S3). Plots of the expression of well-known escapees (*GYG2, ZFX, KDM6A, JPX*) as a function of the number of X chromosomes confirm a consistent expression increase in both NPCs and CMs (Figure S3E, F). In contrast, the inactivated gene *UBE2A* does not show X-copy number dependent expression changes. Note that two cell-type-specific escapees were detected, *PCDH11X* in NPCs and *KLHL4* in CMs, which simply reflects their cell-type specific expression (Figures 1A, B and S3E, F). A notable gene is *ANOS1*, whose expression increases in the presence of a Y in NPCs but not in CMs (Figure 1A, B). Interestingly, the total X expression of non-PAR1 escapees in XX controls represents an average of 125% of that in XY controls, while in XXY the total X expression represents only 115% of that in XY controls, suggesting dampening of X expression in the SCA condition, which was further investigated using allele-specific analyses (Figure S3A, B; Table S3) (see below). Considering X-linked genes classified as subject to XCI, most show similar expression among genotypes (>1.25 log2FC) (Figure S3A, B). Y-linked genes are uniquely expressed in cells bearing a Y chromosome, but surprisingly, their expression differs between XXY and XY and between NPCs and CMs, as further shown below (Figure 1A, B).

**Figure 1.**
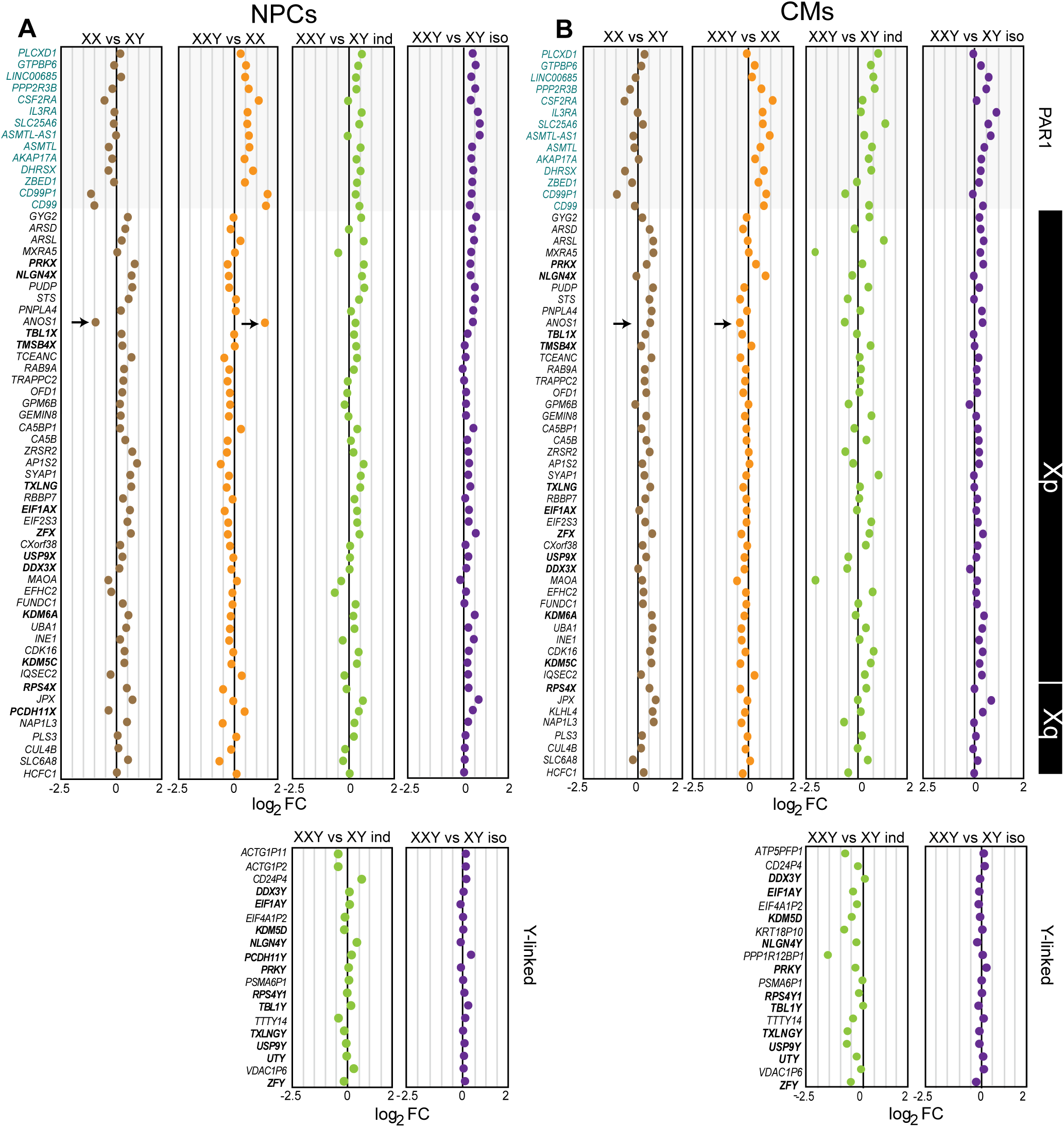
Relative sex-linked gene expression in NPCs and CMs with differential sex chromosome content. **(A, B)** From top to bottom the average log_2_ fold changes of gene expression (CPM) of PAR1 genes, escapees located in Xp and Xq, and Y-linked genes are plotted for the following comparisons: XX vs XY (brown), XXY vs XX (orange), and XXY vs XY either in independent (green) or isogenic lines (purple) of NPCs (**A**) and CMs (**B**). The gene list is on the left and the chromosomal location is noted on the right. The *ANOS1* gene is marked by an arrow, and members of X/Y gene pairs are in bold. *TMSB4Y* is not expressed in NPCs or CMs.

Next, we investigated the effects of SCA on allelic expression of sex-linked genes in XXY. First, we determined changes in expression of X/Y paralogs with copies on both sex chromosomes. Note that the X-linked copy usually escapes XCI and the Y-linked copy either retains functions similar to the X copy or has evolved different functions^[8, 54]^. To measure expression from each sex chromosome, Xa, Xi, and Y, chromosome-specific and allele-specific analyses were done in XX, XY, and XXY clones. The ratios of expression from the Xi [Xi/(Xi+Xa)] and the Xa [Xa/(Xi+Xa)] based on SNP-containing RNA-seq read counts quantify contribution of each allele to the expression of the gene (Figures 2A-D and S4A, B; Tables S4 and S5) (see Methods). In both NPCs and CMs, a decrease in Xi expression (mean decrease of 49% in NPCs and 42% in CMs) is evident in XXY compared to XX for most X/Y gene pairs, which results in lower total X (Xi+Xa) expression in XXY (Figure 2A-D; Table S4). Thus, for X/Y paralogs the dampening of X expression in XXY vs XX specifically results from a decrease in Xi expression.

**Figure 2.**
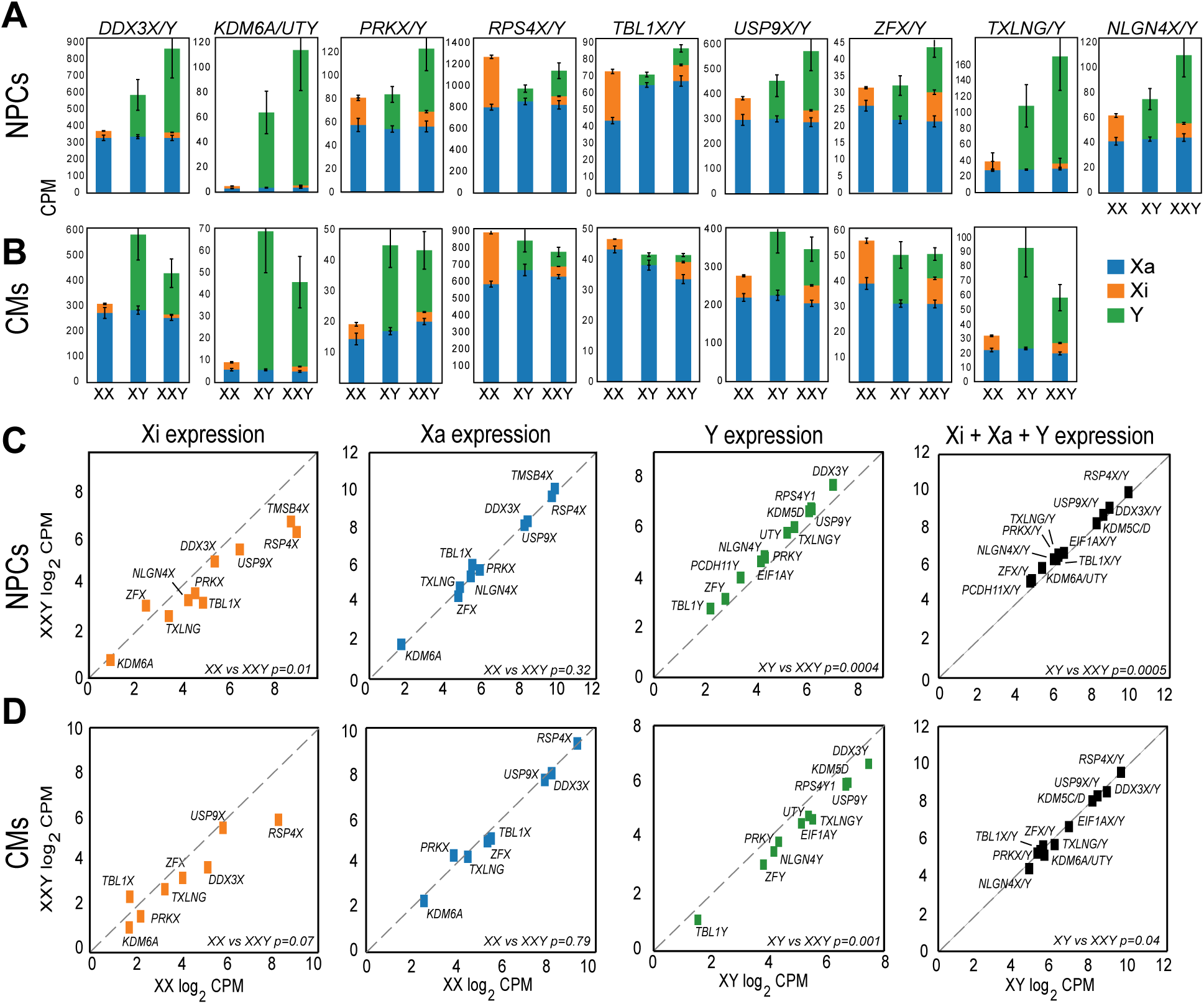
Allelic expression of X/Y genes in NPCs and CMs with a differential sex chromosome content. (**A**, **B**) Stacked bar plots of average CPM expression of homologous X/Y gene pairs (*DDX3X/Y, KDM6A/UTY, PRKX/Y, RPS4X/Y, TBL1X/Y, USP9X/Y, ZFX/Y, TLING/Y, NLGN4X/Y),* in NPCs (**A**) and CMs (**B**) across three genotypes (XX, XY, XXY). Within each bar, expression is partitioned by allelic origin: Xa (blue), Xi (orange), and Y (green). For XX and XXY genotypes, Xa and Xi contributions are estimated using donor-level Xi proportions (mean per genotype), with error bars reflecting the standard error of the mean across clones. For XY and XXY, Y-linked gene expression is included directly from CPM measurements. Only genotypes with at least one donor were included in the calculations (Additional data set 1). (**C**, **D**) Log2 transformed CPM values of expression of X-linked alleles (Xi, Xa), Y-linked copies, and total sex chromosome expression (Xi+Xa+Y) of X/Y gene are plotted in XXY vs XX or vs XY in NPCs (**C**) and CMs (**D**). Xi expression is shown in orange, Xa in blue, Y in green, and total expression (Xi + Xa + Y) in black. Note that there was insufficient data for allelic analysis on *NLGN4X* and *TMSB4X* in CMs, a lack of SNP information for *KDM5C*, *PCDH11X*, and *EIF1AX,* and no expression of *TMSB4Y* in either NPCs or CMs, thus these genes are not included. P-values were determined by Wilcoxon signed rank sum tests. See also Table S4.

Surprisingly, a marked increase in Y expression (ranging from 13-36% higher) is apparent in XXY versus XY NPCs (Figure 2A, C; Table S4). However, this is not the case in CMs where there is lower Y expression (ranging from 25-44% lower) in XXY versus XY (Figure 2B, D; Table S4). In contrast to changes in Xi and Y expression, Xa expression remains similar in NPCs and CMs regardless of the genotype (Figures 2A-D and S4A, B). Our findings indicate that cumulative expression from X and Y copies of each X/Y gene in XXY is modulated by differential cell type-specific changes in Y expression. Indeed, cumulative expression from each X/Y gene (Xa + Xi + Y) in NPCs is highest in XXY compared to XY or XX for most X/Y genes tested (Figure 2A, C and S4A; Table S4), whereas in CMs this cumulative expression is dampened in XXY compared to XY due to lower expression from both the Xi and the Y for most X/Y genes tested (Figure 2B, D and S4B, Table S4). The Y-copy accounts for a high fraction (ranging from 45-95%) of the total expression of some X/Y genes, including *DDX3X/Y*, *KDM6A/UTY* and *TXLNG/Y* (Figures 2A-D). This is consistent with previous studies showing a high contribution of *UTY* to total *KDM6A/UTY* in XY neural cells and in other tissues^[55, 56]^. Note that *UTY* produces more than 200 alternatively spliced transcripts, which may be a confounding factor in determining expression using counts per million (CPM)^[57]^. Indeed, TPM (transcripts per million)-based analyses show similar expression levels between *KDM6A* and *UTY*.

Importantly, our findings of SCA effects on expression from the Xi and the Y are not limited to X/Y paralogs. Indeed, lower expression from the Xi is also observed for escapees without a Y paralog in both NPCs and CMs (Figure 3A, B). These findings indicate widespread expression dampening of supernumerary copies of the X chromosome. Similarly, when considering Y-linked genes that are not members of X/Y genes we also see higher expression in XXY versus XY in NPCs, but lower expression in CMs (Figure 3A, B). We conclude that dampening of gene expression from the Xi occurs in XXY SCA, potentially alleviating deleterious effects of the aneuploidy in both NPCs and CMs. In addition, cell-type specific effects of SCA on Y-linked gene expression are apparent.

**Figure 3:**
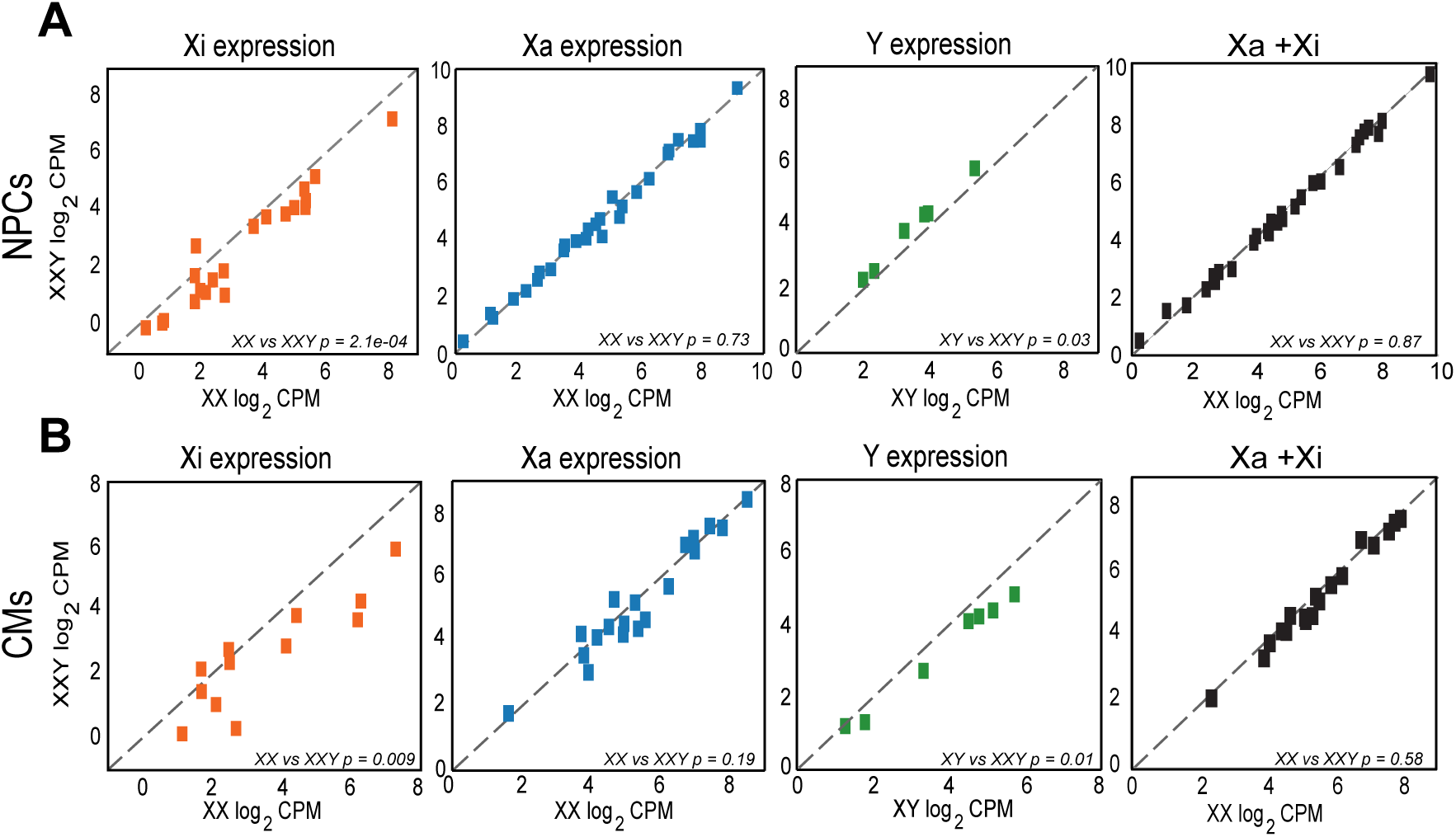
Allelic expression of X-linked escapees and Y-linked genes not included in X/Y pairs in NPCs and CMs with a differential sex chromosome content. (**A**, **B**) Log_2_ transformed CPM values of expression of X-linked escapees (Xa, Xi) and Y-linked genes are plotted in XXY vs XX or XY in NPCs (**A**) and CMs (**B**). Xi expression is shown in orange, Xa in blue, Y in green and total X expression (Xi + Xa) in black. P-values were determined by Wilcoxon signed rank sum tests. See also Table S4.

### Autosomal and overall gene expression in SCA

Differentially expressed autosomal genes were identified using a p-value<0.001 for comparisons between XXY and normal XY and XX controls based on RNA-seq data in isogenic and independent lines of NPCs and CMs (Tables S2 and S6). PCA shows separation between XXY and XY genotypes in comparisons of independent NPC and CM lines. However, considering pairs of isogenic lines, individual clones did not consistently segregate based on their XXY or XY genotype (Figure 4A, B). PCA of independent XXY to XX lines showed genotype-dependent separation in NPCs but limited separation in CMs (Figure 4A, B). The paucity of autosomal DEGs common to isogenic and independent NPCs and CMs is indicative of high variability in autosomal gene expression among clones derived from hiPSCs, which is known to hamper the identification of specific DEGs implicated in SCA (Table S6)^[58]^.

**Figure 4:**
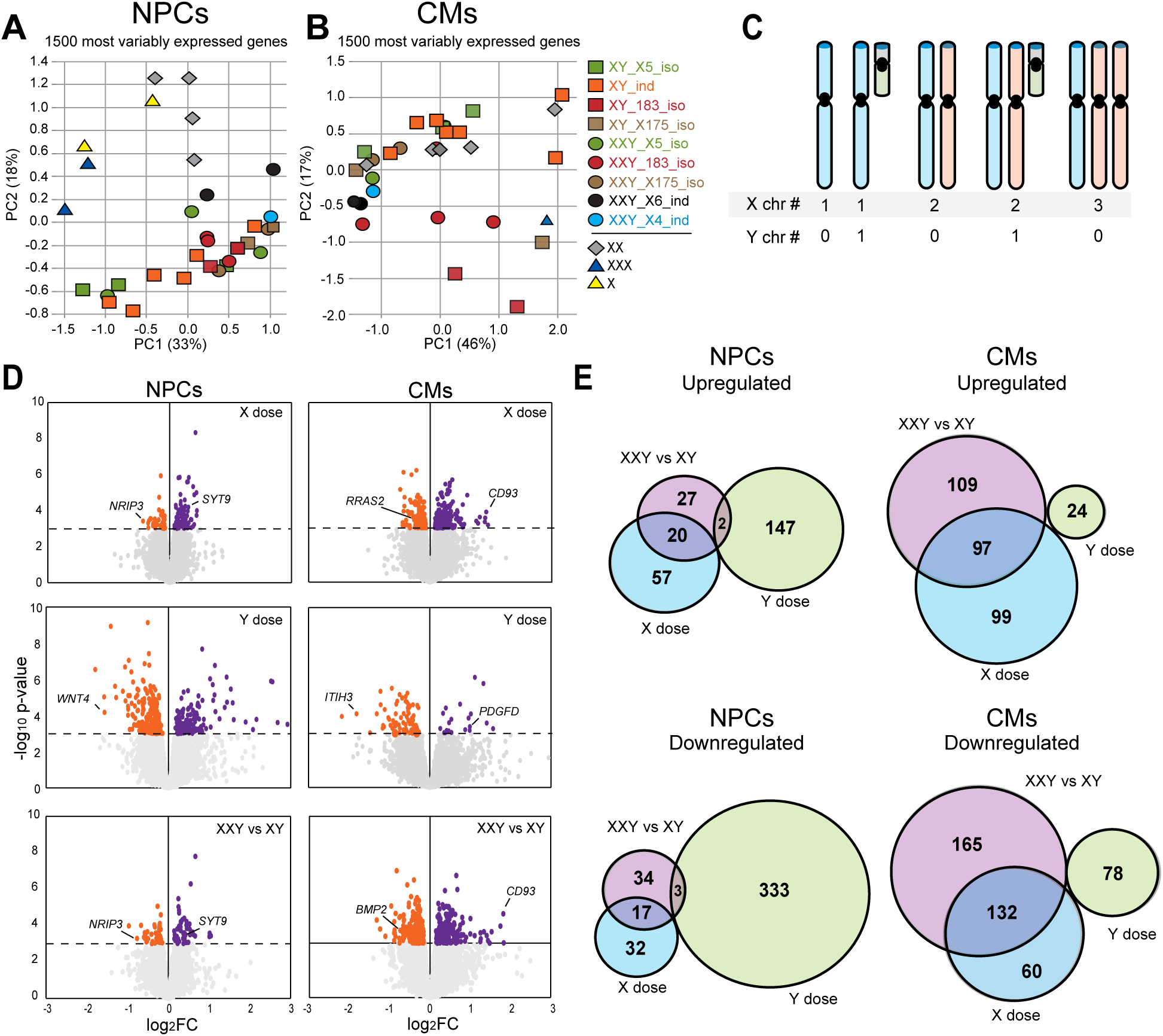
Effects of sex chromosome dosage on autosomal gene expression. (**A**, **B**) PCA plots based on DEG analysis using the 1500 most variable autosomal genes to compare gene expression among clones of NPCs (**A**) and CMs (**B**) with a different number of sex chromosomes. The key shows the list of genotypes with the line names. Isogenic lines are color-coded. Iso: isogenic; ind: independent. The PCA is based on average data for clones within each line. (**C**) Schematic and table of the sex-chromosome copy number series comparisons. The Xa is in blue, Xi in pink, and Y in green. Lines with the same dose of X chromosomes or Y chromosomes were grouped for comparisons. Note that only NPCs include X cells. (**D**) Volcano plots of autosomal genes with expression changes in NPCs and CMs with different sex chromosome doses using a p<0.001 cut off. Purple dots indicate genes with an expression increase and orange dots, genes with an expression decrease. log_2_FC are shown per chromosome X dose (top), per Y dose (middle), and in XXY vs XY cells (bottom). (**E**) Venn diagrams of the number of upregulated and downregulated autosomal genes in XXY vs XY comparisons (purple), in cells with different Y dose (green), and in cells with different X dose (blue) in NPCs and CMs (see also Tables 1 and S7).

Substantial cell-type specific differences were found by comparing the number of autosomal DEGs dependent on the number of Xs (X dose) or the presence of a Y (Y dose) in NPCs and CMs (Figure 4C-E; Tables 1 and S7). Indeed, the Y dose has the greatest influence on autosomal gene expression in NPCs, while the opposite is observed in CMs where the X dose has the greatest influence (Figure 4C-E; Table 1). The effect of the X dose is also apparent when comparing XXY to XY. These findings parallel those based on allelic expression analysis of escapees and Y-linked genes described above, in which higher Y expression is observed in NPCs, in contrast to lower expression in CMs (Figures 2–4). There is no overlap between autosomal DEGs dependent on the Y or X dose, suggesting unique responses to Y and X chromosome dosage in NPCs and CMs (Figure 4D, E; Table S7).

**Table 1:** Number of sex chromosome dosage-sensitive autosomal DEGs.

|  | **Upregulated DEGs** |  |  |
| --- | --- | --- | --- |
| **Cell Type** | **ChrX dose** | **ChrY dose** | **XXY vs XY** |
| NPCs | 77 | 149 | 49 |
| CMs | 196 | 24 | 206 |
|  | **Downregulated DEGs** |  |  |
| **Cell Type** | **ChrX dose** | **ChrY dose** | **XXY vs XY** |
| NPCs | 49 | 336 | 54 |
| CMs | 192 | 78 | 297 |

Next, we investigated the effects of SCA in the regulation of sex-biased autosomal gene expression identified by comparisons of control XX and XY NPCs and CMs. Given the reported subtle differential expression of sex-biased genes^[11]^, all autosomal genes (256 in NPCs, 78 in CMs) with expression differences (p<0.001) were included for analysis (Figure 5A; Table S8). The near 3.3-fold higher number of sex-biased genes in NPCs versus CMs reflects previous findings comparing brain to heart^[11, 59]^. In addition, female-biased genes are especially abundant in NPCs where there is a 4.1 ratio of female- versus male-biased genes, while in CMs this ratio is 2.2 (Figure 5A, D, E; Table S8). Analyses of sex-biased genes in XXY versus XY or XX show that SCA has a marked effect on the expression of sex-biased genes identified in normal controls. Indeed, both NPCs and CMs show a reduction in median fold change in sex-biased autosomal expression in XXY versus XY, regardless of the lines being isogenic or independent (Figure 5B, D, E; Table S8). When comparing XXY to XX, a lesser reduction in sex-biased gene expression is observed in NPCs than in CMs (Figure 5C-E).

**Figure 5:**
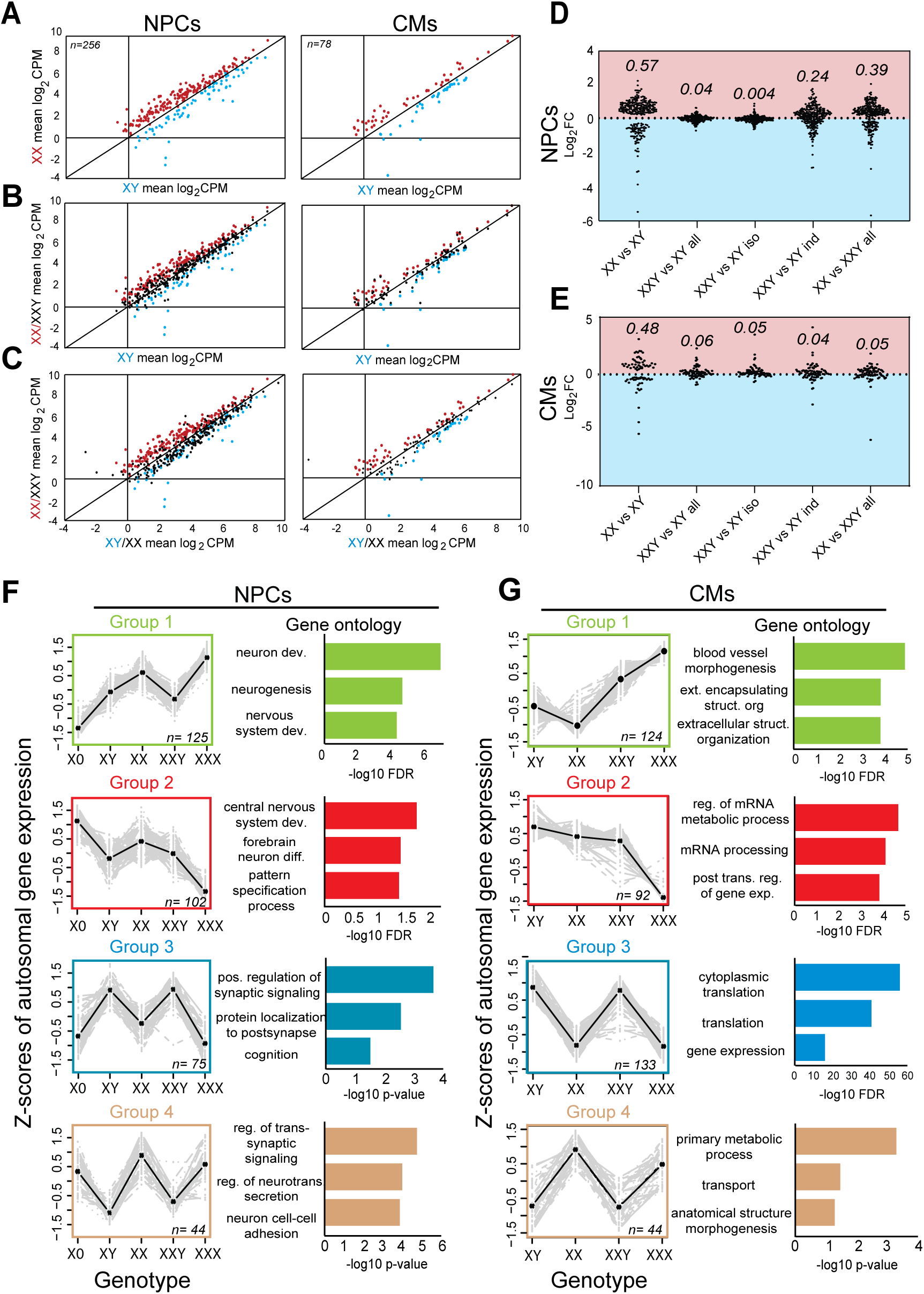
Sex-biased gene expression and K-means analysis of autosomal gene expression changes in cells with differential sex chromosome content. (**A**) Autosomal genes with sex-biased expression were identified based on scatter plots of average CPM values for each gene in normal XY male and XX female NPCs and CMs. Genes with >1.25 log_2_FC at p<0.001 were classified as either female-biased genes (in red) or male-biased genes (in blue). (**B, C**) The sex-biased genes identified in (**A**) were then re-plotted together with comparisons of the same genes in (**B**) XXY vs XY (in black) and (**C**) XXY vs XX (in black). (**D**, **E**) Stack dot plots of female-biased gene expression (light red zone) and male-biased gene expression (light blue zone) illustrate the loss of sex biases in comparisons of XY to XXY, considering all lines (all), isogenic lines (iso) and independent lines (ind). This loss of sex biases is less prominent when comparing XX to XXY lines where biases are better retained in NPCs. (**F**) K-means grouping based on autosomal gene expression in NPCs with different genotypes (X, XX, XY, XXY, XXX). To minimize the effect of high-expressing genes on grouping, log_2_CPM values were converted to z-scores. Graphs of gene expression Z-scores (mean in black and single genes in grey) are shown for groups 1-4, along with the gene ontology. n indicates the number of genes in each group. (**G**) Same analysis as in (**F**) for CMs (XX, XY, XXY, XXX).

To investigate the role of SCA on autosomal gene networks in NPCs and CMs, the top 10% most variably expressed autosomal genes between multiple genotypes (X, XY, XX, XXY, and XXX) were grouped using K-means clustering, which identified four groups with trends dependent of sex chromosomes (Figure 5F, G; Table S9). Group 1 (green) shows increased expression with sex chromosome copy number, group 2 (red) shows the opposite trend, group 3 (blue) shows higher expression in male cells (XY and XXY), and group 4 (tan), higher expression in female cells (X, XX, XXX). Despite identifying these groups in both NPCs and CMs, very few genes are in common between cell types and GO analyses point to gene networks involved in cell-type specific programs.

Identification of gene networks and weighted co-expression network analyses (WGCNA) considering all genes (autosomal and sex-linked) show that among genes whose expression is disrupted by SCA many are associated with cell-type specific functions, e.g. genes associated with neuronal features in NPCs and with metabolism and cardiac function in CMs (Figure 6; Table S10). Clonal variability among derivatives of hiPSCs hampered identification of gene networks with significant changes in SCA. Nonetheless, co-expression networks between independent XX and XY controls include modules (darkred and midnightblue) associated with neurodevelopmental processes and neural cell communication (synapse, cytoplasmic vesicle) in NPCs (Figure 6A), and modules (slateblue1 and coral1) associated with signal transduction, Golgi apparatus, and transferase activity in CMs (Figures 6B). Comparison between all XY to all XXY lines identifies modules (darkorange and brown4) associated with neural cell communication (extracellular space and signal transduction) in NPCs (Figure 6C), and modules (white and cyan) associated with cellular energy production such as mitochondrial and other metabolic processes in CMs (Figure 6D). A similar cell type-specificity in biological processes is observed in comparisons between XX and XXY cells (Figure 6 E, F). Importantly, the co-expression modules in these comparisons are not strongly driven by sex-linked genes as demonstrated by the relatively small percentage of sex-linked genes included. Indeed, we expect about 5% of all genes to be X-linked and 0.7% to be Y-linked. One exception is the presence of most Y-linked genes in the midnightblue and darkgrey modules in NPCs, which is consistent with our findings of a large Y-dosage effect in NPCs whereas no Y-linked genes are included in any module shown in CMs (Table S10).

**Figure 6:**
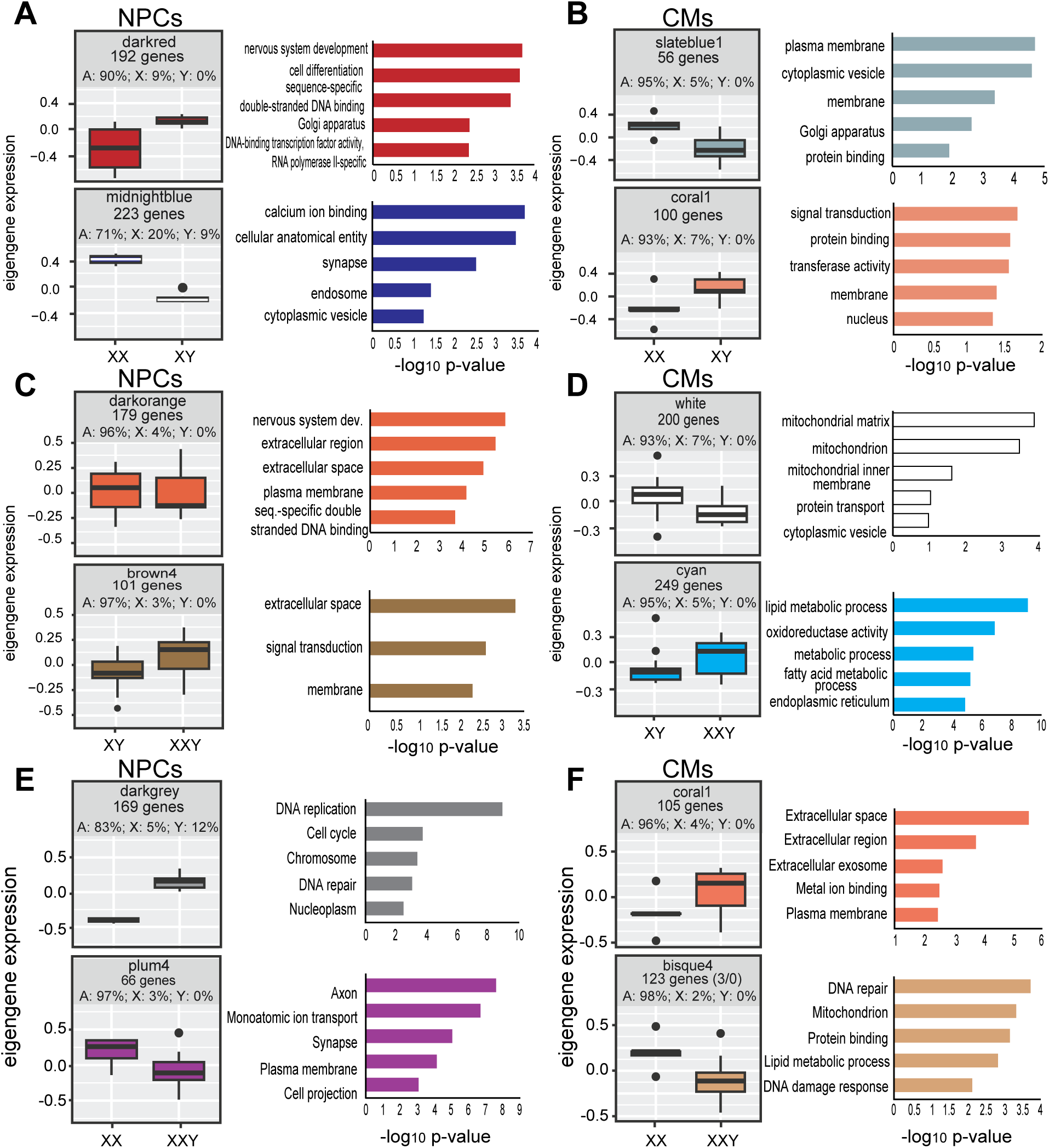
Sex chromosome dosage effects on co-expression modules for all genes. (**A-F**) Box plots of WGCNA analysis shown as eigengene expression and GO terms in three comparisons: (**A, B**) XX vs XY, (**C, D**) XXY vs XY, and (**E, F**) XXY vs XX for NPCs (**A, C, E**) and CMs (**B, D, F**). GO terms are listed for each comparison. For all GO analyses, significant biological processes with >10 genes are shown (FDR <0.05). The total number of genes included in each plot is indicated together with the percentage of autosomal (A), X-linked (X) and Y-linked genes (Y). Note the large number of Y-linked genes in the NPC midnightblue module and in the darkgrey module where the percentage of X-linked is also high. In contrast, there is no evidence of Y-linked genes in CM modules. Based on representation in the human genome, we expect ∼5% X-linked genes and ∼0.7% of Y-linked genes.

### SCA dependent DNA methylation changes

Identification of differentially methylated positions (DMPs) in NPCs and CMs with different sex chromosome complements shows more X-linked hypermethylated DMPs in XX or XXY than XY, consistent with increased XCI-associated promoter DNA methylation (Figure S2C, D). Interestingly, compared to XX lines, XXY lines showed more hypermethylated and fewer hypomethylated X-linked DMPs especially in CMs, which could contribute to dampening of X expression in XXY (Figure S2C, D; Table 2). Consistent with this, we observed higher methylation at CpG islands located near the escapees included in our allelic expression analyses (Figures 2, 3, S4, and 7A, B). No Y-linked promoter CpG island DMPs were found in NPCs or CMs in XXY versus XY comparisons (FDR<0.05), suggesting that Y-linked gene regulation may not depend on DNA methylation. The promoter CpG island of *ANOS1* was hypomethylated in XY and XXY compared to XX NPCs, consistent with NPC-specific Y-dependent *ANOS1* expression (Figures 1A, B and 7C). In contrast to X-linked DMPs, there are more hypomethylated than hypermethylated autosomal DMPs in XXY versus XX in both NPCs and CMs (Table 2). Surprisingly, comparisons between XXY versus XX yielded the highest number of autosomal DMPs in both cell types. Based on MDS clustering using only autosomal genes, isogenic XXY/XY pairs exhibit high inter-variability and low intra-variability in NPCs and CMs, suggesting that members of paired lines have similar epigenetic landscapes (Figure 7D, E). This epigenetic memory may be a feature of mosaic individuals from whom the lines were derived either by cloning or by removing one copy of the X chromosome.

**Figure 7:**
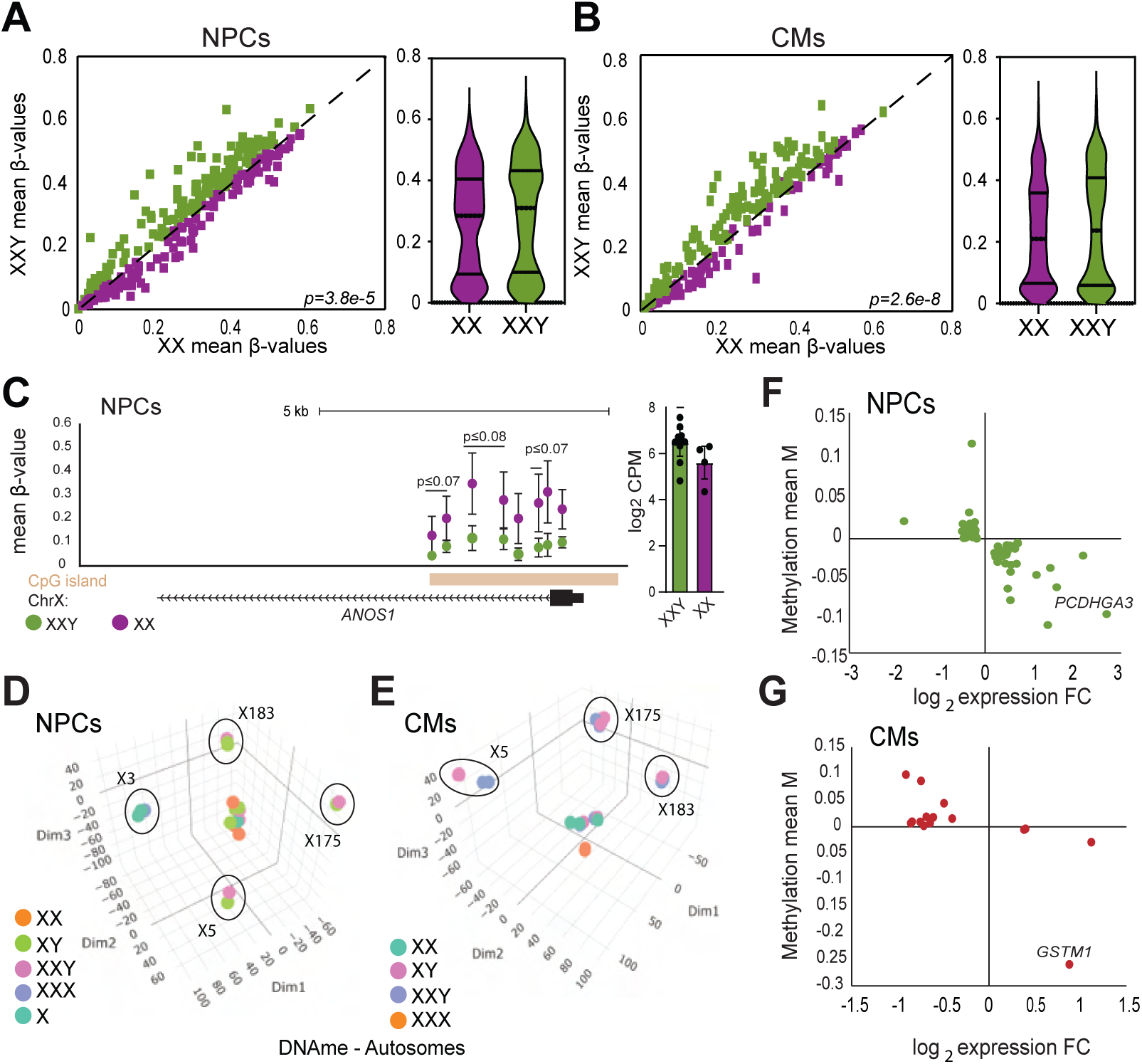
Y chromosome dosage-dependent changes in DNA methylation. **(A, B)** Scatter plots of DNA methylation represented by average β-values show an increase in DNA methylation in XXY vs XX at individual CpGs in islands located near promoters (within 2kb of TSS) of escapees that have decreased expression from the Xi in XXY (see Figures 2 and 3) in NPCs (**A**) and CMs (**B**). CpGs hypermethylated and hypomethylated in XXY vs XX are in green and purple, respectively. Also shown are violin plots illustrating higher β-values in XXY vs XX lines confirming higher methylation levels. (**C**) UCSC genome browser view of DNA methylation changes at the promoter CpG island of *ANOS1* (indicated by a brown bar). Each dot above the CpG island represents the average β-value of methylation of a CpG in XXY lines (green) vs XX lines (purple). The bar graphs on the right show the log_2_CPM expression values for XXY and XX NPCs. (**D**, **E**) 3D MDS plots of DNA methylation in (**D**) NPCs (XX, XY, XXY, XXX and X) and (**E**) CMs (XX, XY, XXY, XXX) using only autosomal M values. The isogenic pairs of lines are circled and labelled X183, X175, X5, and X3 (see Table S1), while the independent lines are not labelled. (**F, G**) Scatter plots showing expression changes anti-correlated to DNA methylation changes at promoter DMRs for autosomal genes that differ between all lines with a Y chromosome (XY and XXY) and lines without a Y chromosome (X, XX, XXX) in NPCs (**F**) and CMs (**G**). Mean of M values representing the level of DNA methylation is on the Y-axis and log_2_FC of gene expression is on the X-axis. *PCDHGA3* in NPCs and *GSTM1* in CMs are highlighted.

**Table 2:** Summary of the number of genes with expression that anti-correlates with DNA methylation at promoter region DMRs.

| Cell type | Gene location | X dose | Y dose |
| --- | --- | --- | --- |
| NPCs | X | 10 | 1 |
|  | Autosomes | 8 | 70 |
|  | Y | 0 | 2 |
|  | **Total** | **18** | **73** |
| CMs | X | 12 | 1 |
|  | Autosomes | 9 | 15 |
|  | Y | 0 | 3 |
|  | **Total** | **21** | **19** |

For each cell type, the number of anti-correlated events between gene expression and DNA methylation located on the X chromosome, the autosomes, and the Y chromosome, are shown with the total as a function of the X chromosome dose or the Y chromosome dose.

Consistent with previous studies, we found few autosomal genes with anti-correlated changes, i.e. low methylation with high expression or high methylation with low expression at differentially methylated regions (DMRs) that overlap the promoter region (1500 bp upstream to 500 bp downstream of the TSS) (Table S11)^[15, 16]^. When comparing cell lines with a Y chromosome to those without, we observed a higher number of anticorrelated genes in NPCs than in CMs, consistent with a greater effect of the Y chromosome on autosomal gene expression in that cell type (Figure 4E; Figure 7F, G; Table S11). In NPCs, genes with anti-correlated changes include genes associated with cognition (e.g. *PCDHGA3*)^[60]^, while in CMs they include genes associated with a risk of myocardial infarction and cardiomyopathy (e.g. *GSTM1*) (Figure 7F, G)^[61]^.

## Discussion

Sex chromosome aneuploidy is associated with disruptions of gene expression and DNA methylation genome-wide resulting in patterns unique to the specific aneuploidy being studied^[9, 15, 16, 31, 62–64]^. However, little is known about the cell type and allele specificity of these effects. We developed disease-relevant cellular models of SCA, namely NPCs and CMs, two cell types representing brain and heart often affected by SCA. Our main focus being XXY associated with Klinefelter syndrome, we derived pairs of isogenic XXY and XY NPCs and CMs, supplemented with independent XXY and normal control XX and XY cell lines. Comparisons between cell lines with different numbers of sex chromosomes revealed allele-specific effects that uncover a novel layer of dosage compensation, which significantly reduces expression of escapees from the inactive X chromosome (Xi) but does not change expression from the active X (Xa) in XXY cell lines. This finding points to a dampening mechanism that may serve to mitigate X chromosome overdosage in XXY NPCs and CMs versus normal cells. We also found major differences in Y-linked gene expression between the two cell types.

Recent studies have proposed that both members of X/Y gene pairs, i.e. the escapee copy on the Xi and the Y-copy, can cross-regulate expression of the Xa in lymphoblastoid cell lines^[17]^. In contrast, our studies of X/Y genes and of other escapees and Y-linked genes in NPCs and CMs point to stable Xa expression together with repression of the Xi copy in XXY, which could be facilitated by the presence of the XCI machinery for X silencing. In fact, many escapees are already partly repressed in normal XX females, with expression levels from the Xi averaging about 30-40% of Xa expression dependent on the cell type^[65–67]^. In XXY, there could be spreading of silencing chromatin marks e.g. H3K27me3, DNA methylation, and chromatin condensation into escape domains. This would lead to a reduction in Xi expression that may represent a positive selection mechanism in SCA. Our analysis of DNA methylation does show more hypermethylated and fewer hypomethylated X-linked differentially methylated positions in XXY than in XX at escapees. Further studies will help clarify the epigenetic mechanisms that control the level of escape in SCA. Together with previous studies showing a unique methylation pattern of the X in Klinefelter, our results support a role for the Y chromosome in regulating DNA methylation on the X^[15, 68, 69]^. Thus, the Y chromosome in XXY compared to XX may trigger dampening of the Xi. There was a larger number of autosomal genes with expression anti-correlated to DNA methylation in NPCs compared to CMs, suggesting a greater dependence of gene expression on DNA methylation levels in NPCs.

A surprising finding in this study is the difference in expression of Y-linked genes in NPCs and CMs, which may reflect the known high expression of Y-linked genes in neural cells^[56]^. As previously reported, many sex-linked genes have high expression in brain and in testis – the so-called brain and balls hypothesis^[70, 71]^. Our analyses show that Y-linked genes are indeed more highly expressed in NPCs than CMs. In XXY NPCs the cumulative expression from the three sex chromosomes is higher than in XX and XY due to higher Y expression and despite the reduction in Xi expression. This is not the case in CMs, suggesting opposite effects in NPCs and CMs. In addition to cell type-specific expression differences at the chromosomal level, we also found differences between cell types at the single gene level: for example, in NPCs but not in CMs the known escapee, *ANOS1*, shows gene expression and methylation patterns driven by the presence of the Y chromosome. *ANOS1* causes Kallmann Syndrome, a highly male-biased disorder characterized by distinct facial features, delayed puberty, and hearing loss^[72, 73]^.

Sex chromosome aneuploidy disrupts sex biases in autosomal gene expression in a cell-specific manner, which is not surprising given that there are almost three times as many autosomal genes with sex-bias in NPCs compared to CMs, consistent with the brain’s sexualized transcriptome^[74]^. There is a predominance of female-biased genes in both cell types, and a marked reduction in sex-biased expression occurs in XXY versus XY in both NPCs and CMs, reflecting widespread effects of SCA^[18, 62–64, 75]^. We found that the sex-bias effect of an added Y-chromosome to XX NPCs is much less than that caused by an added X chromosome to XY cells, consistent with the large gene content of the X compared to the Y^[2]^. Similar effects have been reported in other human cell types^[9]^. However, in CMs the effects of added Y or X are similar, suggesting that sex-biased gene expression is sensitive to both X and Y chromosome dosage in this cell type.

Most of the cell-type specific grouping of autosomal DEGs dependent on sex chromosome dosage may play roles in the underlying brain- or heart-related phenotypes associated with SCA^[2, 76, 77]^. For example, among the Y-dependent autosomal genes found in NPCs *WNT4* and the protocadherin genes have important roles in neurogenesis and brain development^[78, 79]^. Among genes downregulated with increasing X chromosome copy number in NPCs, *NRIP3* is highly expressed in brain and testis and may be associated with hypogonadism, seen in XXY^[80, 81]^. Synaptotagmin 9 (*SYT9*), a gene upregulated with increasing X or sex chromosome copy number in NPCs, is a Ca2+ sensor expressed in brain and the pituitary and implicated in reproductive hormone secretion, which is compromised in XXY^[82]^. *SYT9* is part of a large family of membrane trafficking proteins that trigger synaptic vesicle exocytosis in striatal neurons with crucial roles in motor control, a mechanism often disrupted in Klinefelter individuals^[83, 84]^. Genes with co-expression changes based on X-dosage or Y-presence are linked to communication between neural cells, which is in line with previous findings in mouse and human that describe signaling differences between normal males and females^[84, 85]^. Autosomal DEGs identified in CMs include several genes associated with critical heart functions, e.g. *CD93* that responds to X copy number and *ITIH3* and *PDGFD*, both responding to the presence of the Y^[86–88]^. *KRT8* and *KRT18* two keratin genes included in the Y-copy-dependent group 3 identified by K-means analysis in CMs have a cardioprotective role in mice by maintaining intercalated disc structure and mitochondrial health in a desmin-deficient mouse model^[89]^. A subset of kinesin genes identified with higher expression in XXY than XY CMs includes *KIF20A*, a gene mutated in individuals with congenital heart defects similar to those seen in TS^[90]^. In addition, *RRAS2* and *BMP2*, two genes whose expression decreases with increasing X or sex chromosome number, have been associated with cardiac functions: mutations in *RRAS2* cause Noonan Syndrome associated with cardiac abnormalities, while *BMP2* is associated with increased aortic valvular calcification, a condition more prevalent in Turner syndrome compared to XX^[91–93]^.

## Conclusions

Our study based on cellular models of brain and heart in SCA has uncovered evidence of a new mechanism of dosage compensation in an aneuploidy condition. Indeed, allele-specific dampening of expression from genes expressed from the inactive X chromosome was observed in both NPCs and CMs with a XXY genotype compared to a XX genotype. Although the epigenetic mechanisms of this adjustment in expression of X-linked genes remain to be identified, the presence of effectors of gene silencing on the inactive X may facilitate the beneficial adjustment in gene expression in SCA. Variable levels of dampening could potentially explain variable expressivity of Klinefelter syndrome. We also found striking differences between cell types in levels of Y-linked genes expression and in responses of autosomal genes to the presence of a Y chromosome. These unexpected Y effects will need to be defined in other cell types. Our analyses further identified a number of cell-specific genes with altered expression in XXY, which will help understanding of this condition.

## Supporting information

Addtional dataset 1

## Resource availability

Gene expression and DNA methylation sequencing data used in this study has been deposited in the NCBI GEO database with accession numbers GSE324828 and GSE324692, respectively. All other data and the scripts used for the analyses that support the findings of this study are available from the corresponding authors upon reasonable request.

## Author contributions

Conceptualization: C.M.D., J.B.B., X.D., G.N.F; Investigation: J.B.B., G.N.F., A.M., E.S., X.D., C.G., A.S., M.T., C.M.D; Material acquisition: D.V.D., A.S., C.H.G., C.M.D; Formal data analysis: W.M, T.B, J.M., H.F., E.S., G.L., R.Z., W.S.N; Methodology: G.N.F., J.E.Y., C.E.M., Y.L., W.S.N; Data Curation: T.B., J.M.; Writing and Visualization: J.B.B., G.N.F., T.B., J.M.; X.D. and C.M.D.; Supervision, J.B., X.D., G.N.F., W.S.N., C.M.D.; Funding Acquisition: C.M.D, J.E.Y, J.B.B., W.S.N., W.M.

## Acknowledgments

This work was supported by NIH grants GM113943 (C.M.D.), GM131745 (C.M.D.), HG011586 (C.M.D., W.S.N., X.D.), MH105768 (J.B.B.), AG073918 (C.M.D. and J.E.Y.), and NSF DBI175317 (W.M.). The funders had no role in the design of the study and collection, analysis, and interpretation of data and in writing the manuscript. We would like to thank Dr. David Russell for providing technical expertise in selection methods for removal of an X chromosome to generate isogenic hiPSCs. We would also like to thank Drs. Lil Pabon and Refugio Martinez for their technical help in differentiating CMs and NPCs, respectively. A special thank you to Dr. Julie Mathieu, Chris Cavanaugh and Jenn Hesson at the Tom and Sue Ellison Stem Cell Core at the University of Washington Institute for Stem Cell and Regenerative Medicine for their technical expertise and assistance especially with mosaic hiPSC reprogramming.

## Supplemental Information

Document S1: Supplemental Figures S1-S4

Supplemental table S1: Summary of cell lines

Supplemental table S2: Summary of comparisons of gene expression between genotypes

Supplemental table S3: Escape gene expression as a function of the number of X chromosomes

Supplemental table S4: Allelic expression of escapees in NPCs and CMs

Supplemental table S5: Correlation of allele-specific expression of X-linked SNPs between clone pairs from the same donor

Supplemental table S6: Autosomal DEGs in comparisons between genotypes

Supplemental table S7: Sex chromosome dose dependent autosomal gene expression changes

Supplemental table S8: Sex-biased autosomal genes in NPCs and CMs

Supplemental table S9: K-means clustering of the top 10% most variable autosomal genes in NPCs and CMs

Supplemental table S10: Genes included in highlighted WGCNA modules

Supplemental table S11: Autosomal genes with anti-correlated changes in expression and DNA methylation at nearby promoter DMRs

Data S1: Additional data set 1

