## Supplemental figures and legends for "Cell-type specific allelic dampening of sex-linked genes in sex chromosome aneuploidy"

S1

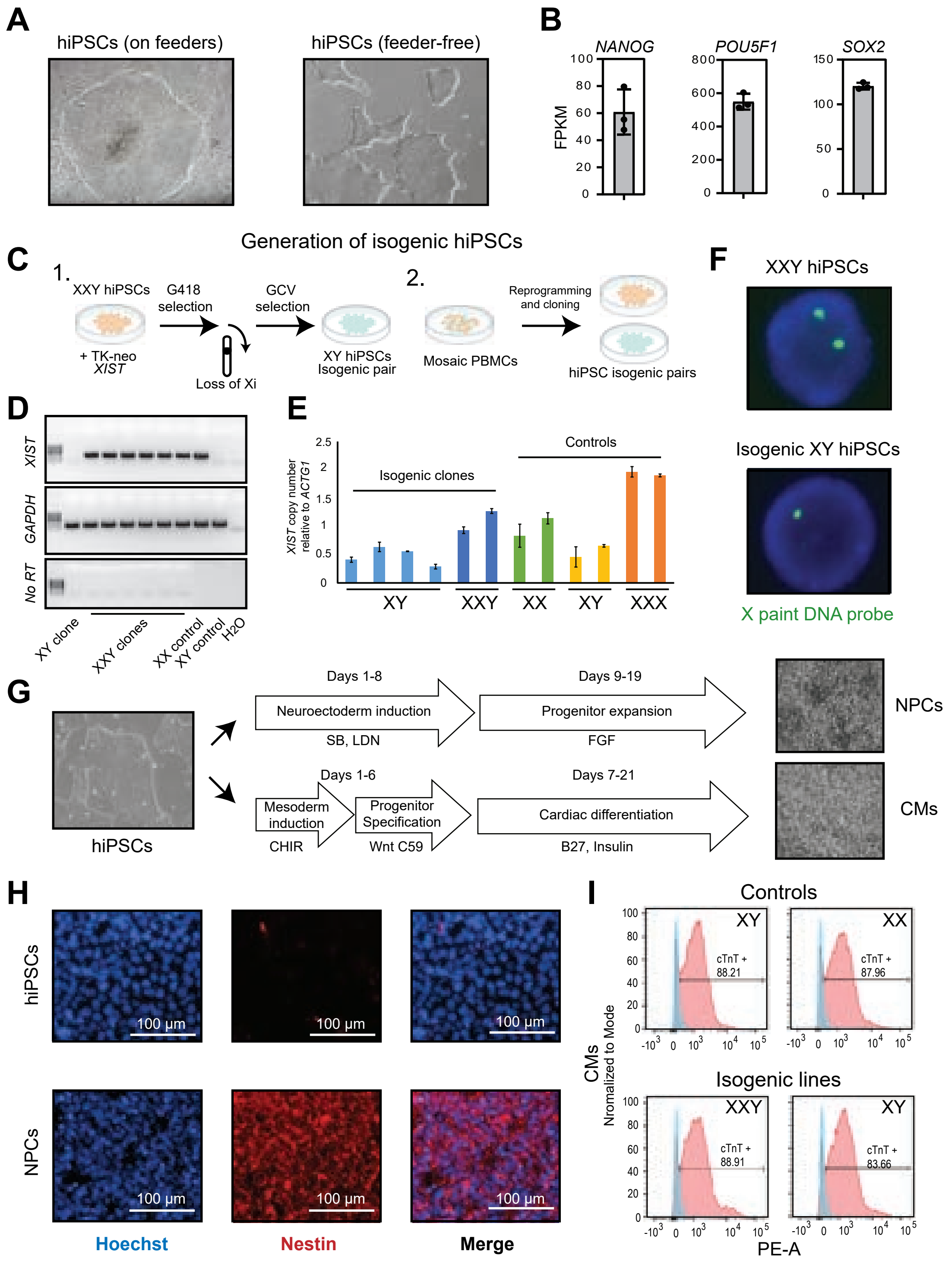

A
hiPSCs (on feeders)
hiPSCs (feeder-free)
B
NANOG
POU5F1
SOX2
FPKM
C
Generation of isogenic hiPSCs
1.
XXY hiPSCs
+ TK-neo XIST
G418 selection
Loss of Xi
GCV selection
XY hiPSCs Isogenic pair
2.
Mosaic PBMCs
Reprogramming and cloning
hiPSC isogenic pairs
D
XIST
GAPDH
No RT
XY clone
XXY clones
XX control
XY control
H2O
E
XIST copy number relative to ACTG1
Isogenic clones
Controls
XY
XXY
XX
XY
XXX
F
XXY hiPSCs
Isogenic XY hiPSCs
X paint DNA probe
G
hiPSCs
Days 1-8
Neuroectoderm induction
SB, LDN
Days 9-19
Progenitor expansion
FGF
NPCs
Days 1-6
Mesoderm induction
CHIR
Progenitor Specification
Wnt C59
Days 7-21
Cardiac differentiation
B27, Insulin
CMs
H
hiPSCs
NPCs
100 µm
Hoechst
Nestin
Merge
I
Controls
XY
cTnT + 88.21
XX
cTnT + 87.96
Isogenic lines
XXY
cTnT + 88.91
XY
cTnT + 83.66
CMs
Nromalized to Mode
PE-A

S2

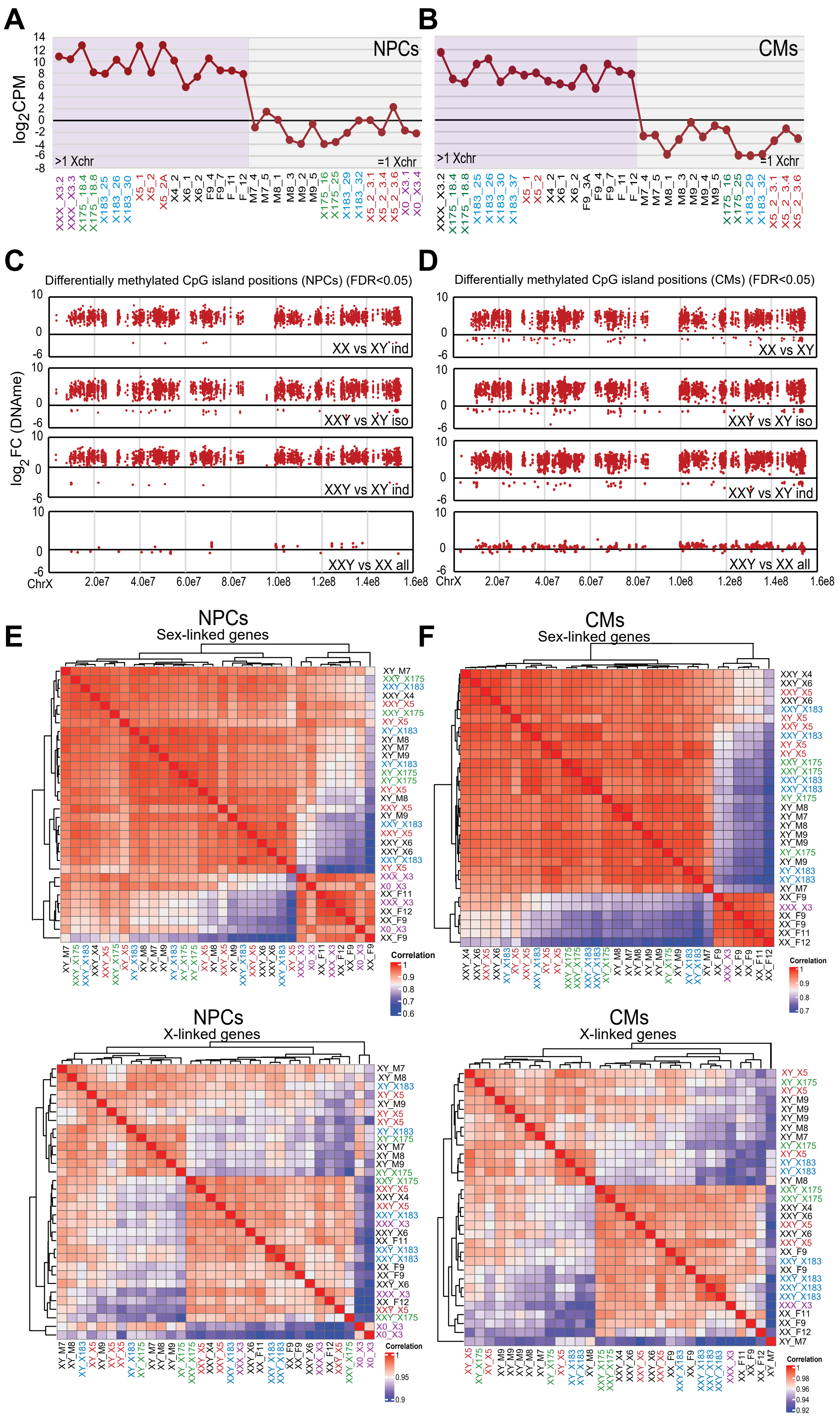

A
B
NPCs
CMs
log2CPM
>1 Xchr
=1 Xchr
C
D
Differentially methylated CpG island positions (NPCs) (FDR<0.05)
Differentially methylated CpG island positions (CMs) (FDR<0.05)
log2FC (DNAme)
XX vs XY ind
XX vs XY
XXY vs XY iso
XXY vs XY ind
XXY vs XX all
ChrX
E
F
NPCs
Sex-linked genes
CMs
Sex-linked genes
NPCs
X-linked genes
CMs
X-linked genes
Correlation

S3

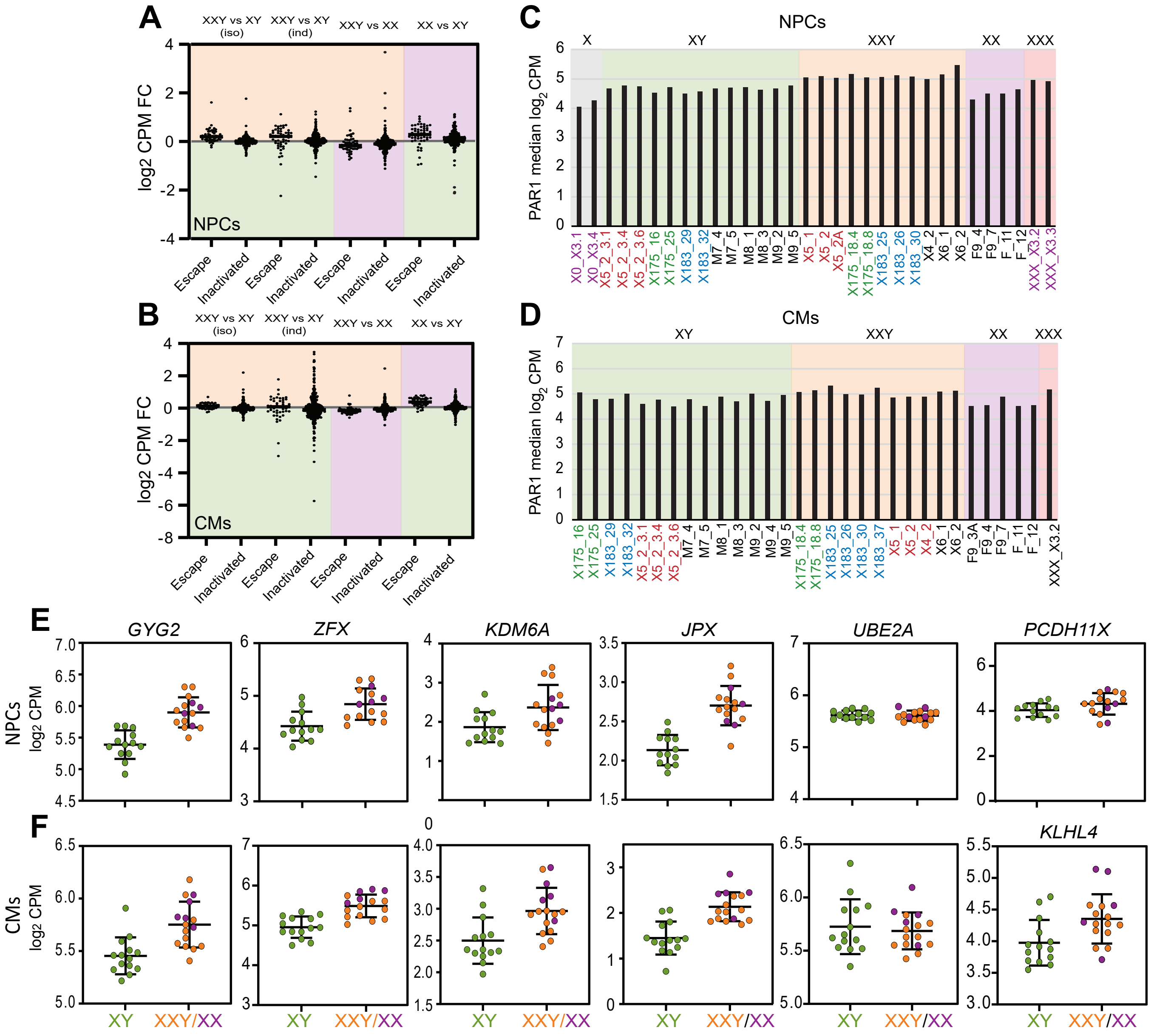
A
XXY vs XY (iso)
XXY vs XY (ind)
XXY vs XX
XX vs XY
log2 CPM FC
NPCs
Escape
Inactivated
B
CMs
C
NPCs
X
XY
XXY
XX
XXX
PAR1 median log2 CPM
D
CMs
E
GYG2
ZFX
KDM6A
JPX
UBE2A
PCDH11X
NPCs log2 CPM
F
KLHL4
CMs log2 CPM
XY
XXY/XX

S4

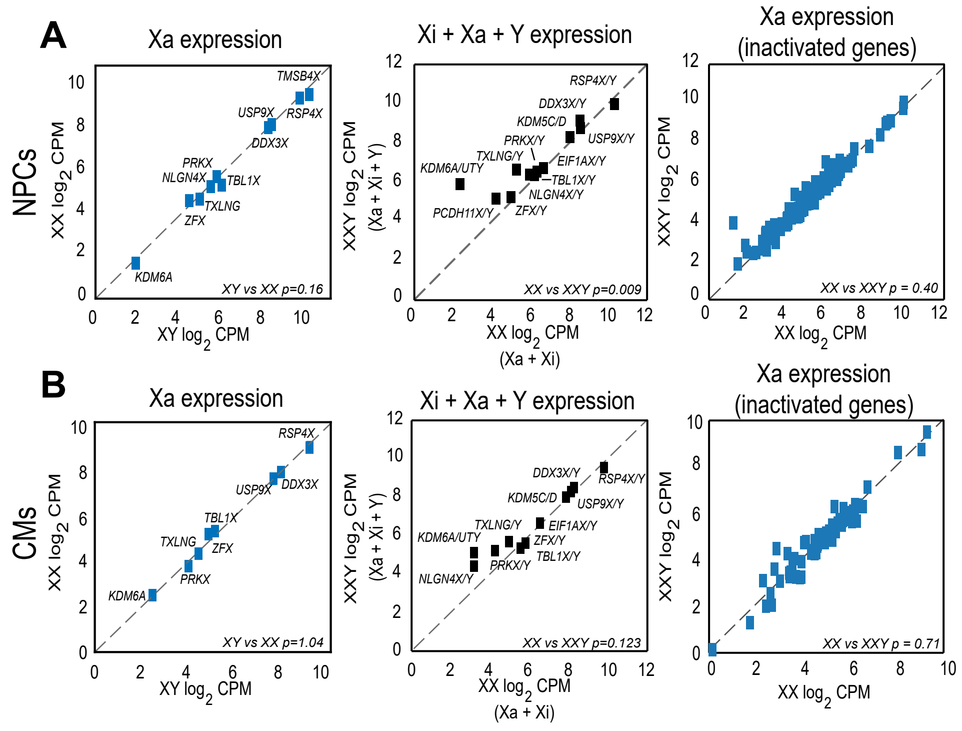
A
NPCs
Xa expression
XX log2 CPM
XY log2 CPM
TMSB4X
USP9X
RSP4X
DDX3X
PRKX
NLGN4X
TBL1X
TXLNG
ZFX
KDM6A
XY vs XX p=0.16
Xi + Xa + Y expression
XXY log2 CPM
(Xa + Xi + Y)
XX log2 CPM
(Xa + Xi)
RSP4X/Y
DDX3X/Y
KDM5C/D
USP9X/Y
PRKX/Y
TXLNG/Y
EIF1AX/Y
KDM6A/UTY
TBL1X/Y
NLGN4X/Y
PCDH11X/Y
ZFX/Y
XX vs XXY p=0.009
Xa expression
(inactivated genes)
XXY log2 CPM
XX log2 CPM
XX vs XXY p = 0.40
B
CMs
Xa expression
XX log2 CPM
XY log2 CPM
RSP4X
USP9X
DDX3X
TBL1X
TXLNG
ZFX
PRKX
KDM6A
XY vs XX p=1.04
Xi + Xa + Y expression
XXY log2 CPM
(Xa + Xi + Y)
XX log2 CPM
(Xa + Xi)
DDX3X/Y
RSP4X/Y
KDM5C/D
USP9X/Y
TXLNG/Y
EIF1AX/Y
KDM6A/UTY
ZFX/Y
TBL1X/Y
NLGN4X/Y
PRKX/Y
XX vs XXY p=0.123
Xa expression
(inactivated genes)
XXY log2 CPM
XX log2 CPM
XX vs XXY p = 0.71

***Figure S1: Generation of isogenic and independently derived hiPSCs, NPCs and CMs with a different number of sex chromosomes.*** (**A**) Representative images of hiPSCs derived from patients with X chromosome aneuploidy and control individuals. hiPSCs show defined colonies with smooth edges when cultured on inactivated MEF feeders as well as in feeder-free conditions. Bright-field mages taken at 5x magnification. (**B**) Bar graphs showing expression in FPKM of *NANOG, POU5F1* and *SOX2* in reprogrammed hiPSCs, confirming the pluripotent state of the cells. (**C**) Schematic of the methods used to generate isogenic pairs of hiPSCs derived by targeting the *XIST* locus by a dual selection TK-neo cassette and selecting for loss of the Xi in XXY hiPSCs from patient amniocytes (1) or by cloning cell lines from hiPSCs derived from blood of mosaic patients (2). (**D**) Representative RT-PCR analysis of *XIST* expression used to screen XXY/XY isogenic hiPSC clones along with XX and XY controls. *GAPDH* serves as a loading control, no-RT - as a no reverse transciptase control, and H2O - as a no-template control. An XY isogenic clone shows loss of XIST expression, consistent with the loss of the Xi. (**E**) Representative *XIST* copy number qPCR analysis used to verify genotypes of XXY/XY isogenic hiPSC clones along with XX, XY, and XXX controls. *ACTG1*, a single copy gene, is used for normalization. Cells with more than one X chromosome show higher copy number than those with one X with XXX showing the highest. (**F**) Examples of nuclei from the X5 hiPSC line (XXY) and its isogenic XY line following DNA-FISH using a fluorescein-labelled X chromosome specific probe, show two green signals in XXY and one green signal in XY. High resolution fluorescent mages taken at 100x magnification. (**G**) Schematic illustrating the differentiation protocols for hiPSC differentiation to NPCs (top) and CMs (bottom). Bright-field images of hiPSCs, NPCs and CMs taken at 10x magnification. (**H**) hiPSCs and NPCs stained by a nestin antibody show staining (red) specifically in NPCs. Nuclei staining using Hoechst in blue. High resolution wide-field fluorescent images taken at 20x magnification. (**I**) Flow analysis of CMs following staining for cardiac troponin T (cTnT) antibody or control isotype IgG1 show specific cTnT+ staining in CMs from control XY and XX and from isogenic XXY/XY lines (red) compared to IgG (blue).

***Figure S2: XCI stability and sex-linked gene expression in differentiated NPCs and CMs.*** (**A, B**) $Log_2$ CPM values for *XIST* expression in each NPC (**A**) and CM (**B**) clone analyzed. *XIST* expression is high in cell lines with more than one X chromosome and is not expressed or is very low in lines with one. Clones are labelled with genotype and color coded by isogenic pair. (**C, D**) DNA methylation fold changes ($log_2FC$) along the human X chromosome at CpG islands in comparisons of XX vs XY (independent), XXY vs XY (isogenic), XXY vs XY (independent) and XXY vs XX (all) NPCs (**C**) and CMs (**D**) (FDR < 0.05). Each dot represents a CpG. Hypermethylation of CpGs occurs in cell lines with two X chromosomes due to XCI which is maintained during differentiation. (**E, F**) Hierarchical clustering heatmaps using X- and Y-linked genes (top) show clear separation of clones based on the presence of the Y chromosome, while hierarchical clustering using X-linked genes only (bottom) shows overall clone separation based on the number of X chromosomes in both NPCs (**E**) and CMs (**F**). Clones are labelled with genotype and color coded by isogenic pairs.

***Figure S3: PAR1, non-PAR escapees, and genes subject to XCI in NPCs and CMs with different numbers of sex chromosomes.*** (**A, B**) Stack dot plots of expression of non-PAR1 escapees and inactivated X-linked genes expression in comparisons between NPCs (**A**) and CMs (**B**) with a differential sex chromosome content. Genotypes are color-coded: XX in purple, XY in green and XXY in light orange. Iso: isogenic, ind: independent, all: all clones. $Log_2$ CPM FC is shown. (**C, D**) Median $log_2$ CPM values for PAR1 genes in NPCs (**C**) and CMs (**D**) clones stratified by genotype (X, XY, XXY, XX, XXX). Clone labels are color-coded for isogenic pairs. (**E, F**) $Log_2$ CPM values of known non-PAR1 escapees (*GYG2, ZFX, KDM6A,* and *JPX*), and the inactivated gene *UBE2A*, in NPCs (**E**) and CMs (**F**). The NPC-specific escapee *PCDH11X* and the CM-specific escapee *KLHL4* are also shown. Each dot represents expression in an individual cell line. Genotypes are color-coded: XY (green) XX (purple) and XXY (light orange).

***Figure S4: Allelic expression of X-linked genes in NPCs and CMs with a differential sex chromosome content.*** (**A, B**) Log2 transformed CPM plots show Xa expression values of escapees with a Y paralog plotted in XY vs XX (left), total sex chromosome expression (Xa +Xi + Y) of X/Y genes plotted in XX vs XXY (middle), and Xa expression of all inactivated genes in XX vs XXY (right) in NPCs (**A**) and CMs (**B**). P-values were determined by Wilcoxon signed rank sum tests. See also Table S4.
